# Scaling between stomatal size and density in forest plants

**DOI:** 10.1101/2021.04.25.441252

**Authors:** Congcong Liu, Christopher D. Muir, Ying Li, Li Xu, Mingxu Li, Jiahui Zhang, Hugo Jan de Boer, Lawren Sack, Xingguo Han, Guirui Yu, Nianpeng He

## Abstract

The size and density of stomatal pores limit the maximum rate of leaf carbon gain and water loss (*g*_max_) in land plants. The limits of *g*_max_ due to anatomy, and its constraint by the negative correlation of stomatal size and density at broad phylogenetic scales, has been unclear and controversial. The prevailing hypothesis posits that adaptation to higher *g*_max_ is typically constrained by geometry and/or an economic need to reduce the allocation of epidermal area to stomata (stomatal-area minimization), and this would require the evolution of greater numbers of smaller stomata. Another view, supported by the data, is that across plant diversity, epidermal area allocated to guard cells versus other cells can be optimized without major trade-offs, and higher *g*_max_ would typically be achieved with a higher allocation of epidermal area to stomata (stomatal-area increase). We tested these hypotheses by comparing their predictions for the structure of the covariance of stomatal size and density across species, applying macroevolutionary models and phylogenetic regression to data for 2408 species of angiosperms, gymnosperms, and ferns from forests worldwide. The observed stomatal size-density scaling and covariance supported the stomatal-area increase hypothesis for high *g*_max_. A higher *g*_max_ involves construction costs and maintenance costs that should be considered in models assessing optimal stomatal conductance for predictions of water use, photosynthesis, and water-use efficiency as influences on crop productivity or in Earth System models.

---

Stomatal pores are critical determinants of the function of plants and the composition of the atmosphere (1). The stomatal conductance to diffusion of water vapor and CO_2_ (*g*_s_) influences a broad spectrum of ecological processes at leaf, community, and ecosystem scales, including photosynthesis, net primary production, and water use efficiency (2, 3). Theoretically, stomata can regulate *g*_s_ through evolutionary or plastic shifts in stomatal size or numbers (4) or through short-term stomatal aperture changes (5). The *g*_s_, and its typical operational value (*g*_op_), can thus vary from near zero with stomata fully closed and *g*_max_ with stomata fully open. The *g*_max_ is a fundamental anatomical constraint, and across species measured under controlled conditions, *g*_op_ and *g*_max_ are correlated (6, 7). Because of their importance in controlling leaf water and CO_2_ fluxes, stomatal anatomy can provide critical information in global vegetation and crop models (8-11) toward the current grand challenge of understanding how crops and forest trees are optimized for carbon gain versus water use. Yet, there has been substantial debate about the anatomical underpinnings of the evolution of higher *g*_max_, and its associated costs.

The *g*_max_ is a mathematic function of underlying anatomical traits stomatal density (*D*_s_, number of pores per unit epidermal area) and size (*A*_s_, area of guard cells surrounding each pore). Indeed, these traits are widely used to study the adaptation and competition of plants because they are reliable indicators of *g*_max_(12-18). Further an inverse relationship between *A*_s_ and *D*_s_ across diverse plant species has been recognized since 1865 (19). A prevalent view in the literature established by Franks and Beerling (20) is that the negative *A*_s_ and *D*_s_ relationship and the cost of stomatal area place a strong constraint on the evolution of *g*_max_. According to an early version of the “stomatal area minimization hypothesis” a packing limit geometry constrains *g*_max_, because the total fraction of epidermal area allocated to stomata (*f*_*s*_) cannot exceed unity:

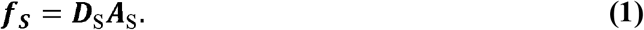

This, in turn generates the negative *A*_s_ and *D*_s_ relationship such that the evolution of larger numbers of stomata would necessitate reduction in their size. Thus, higher *g*_max_ can only be achieved by the evolution of larger numbers of smaller stomata. This early packing geometry argument was rendered moot given observations that for functional leaves, maximum *f*_*s*_ is usually far lower than unity (33.6% in our data; solid line in Fig. 1a). First, because stomata need to be spaced out by epidermal cells to open and close properly (21), and second, because the development of higher *D*_s_ can occur through the increased differentiation of epidermal cells into stomata (i.e., achieving higher stomatal differentiation rate, or stomatal index (22, 23), such that stomatal numbers can be independent of sizes. Yet the stomatal area minimization hypotheses for the evolution of higher *g*_max_ and its association with the negative *A*_s_ and *D*_s_ relationship was reached by a different argument: that to minimize stomatal construction and maintenance costs (24), plants evolving higher *g*_max_ must do so with a reduced *f*_*s*_, and this maximization of *g*_max_ relative to *f*_*s*_ would in turn generate the negative *A*_s_ and *D*_s_ relationship (20).

**Fig. 1.**
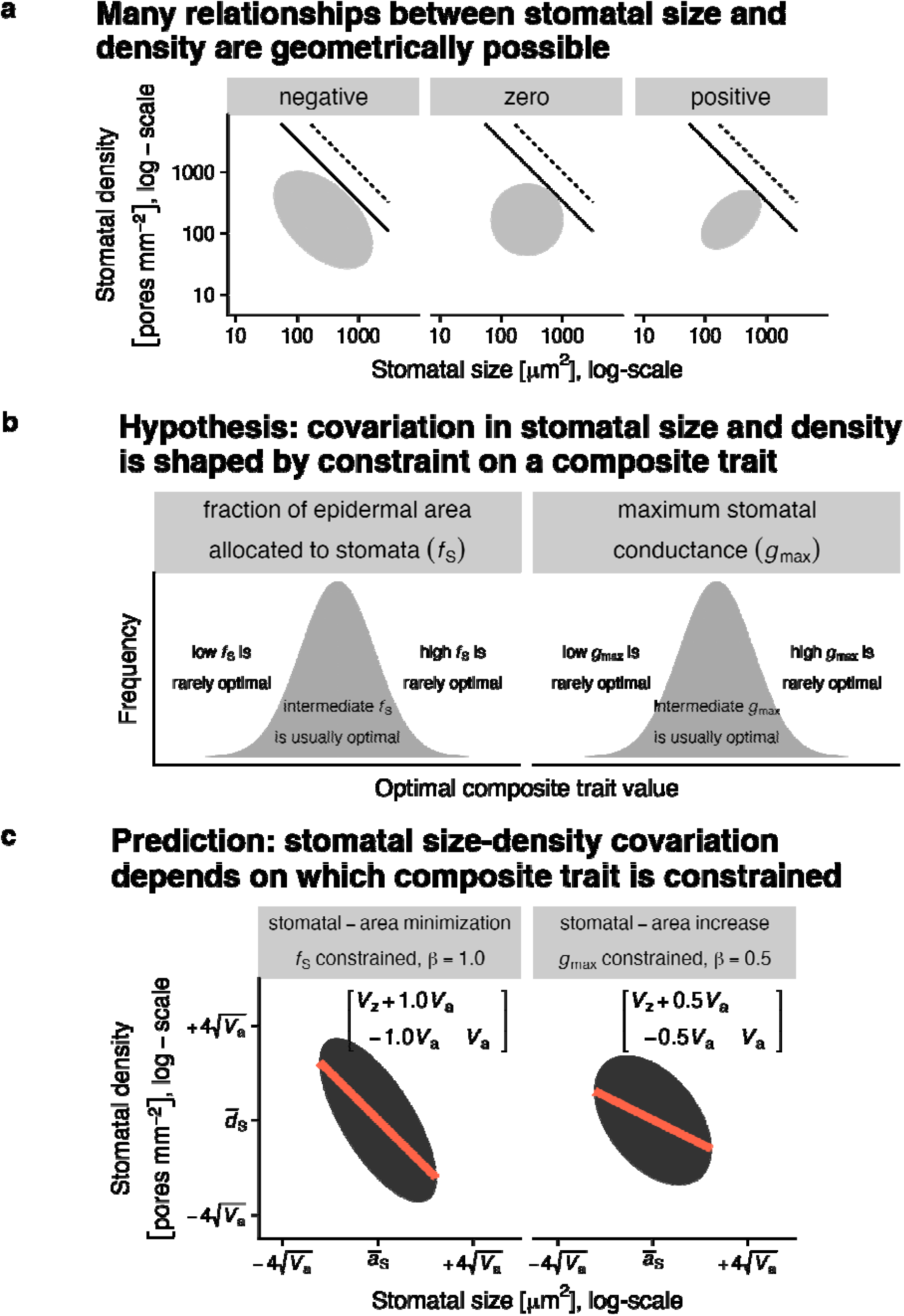
Competing hypotheses for stomatal size-density scaling make different predictions about the trait covariance structure. Maximum stomatal conductance (*g*_max_) and the fraction of epidermal area allocated to stomata (*f*_S_) are composite traits determined by stomatal density and size. On a log-scale, they are the sum of log-stomatal density (*d*_S_) and log-stomatal size (*a*_S_) times a scaling exponent (β), 0.5 for *g*_max_ and 1.0 for *f*_S_ (see Methods). **a**. Many scaling relationships between stomatal size and density are possible as long as *f*_S_ does not exceed 1 (dashed line) or more realistically a value less than 1 to allow space between stomata (solid line, *f*_S_ = 0.34, the maximum value in our data set). The grey ellipses represent different possible scaling relationships with the same mean trait values in our data set (*Ā*_S_= 263 *μ*m^2^, 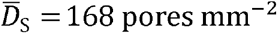). These are 95% quantile of covariance ellipses for a bivariate normal with trait correlations of -0.5, 0, and 0.5 and trait variances of 0.75, 0.55, and 0.45 for ‘negative’, ‘zero’, and ‘positive’ relationships, respectively. **b**. We hypothesized that size-density scaling is determined by constraint on either *g*_max_ (stomatal-area increase; left panel) or *f*_S_ (stomatal-area minimization; right panel). Under either hypothesis, the optimal composite trait varies but extreme values of the composite trait are rarely optimal. **c**. Both hypotheses predict negative size-density scaling but with different covariance relationships. If the interspecific means 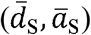 and variances (*V*_*d*_, *V*_*a*_) of stomatal density and size, respectively, are measured, the covariance between them (*V*_*d,a*_) is equal to -β*V*_*a*_. Under the stomatal-area increase (left panel) and stomatal-area minimization (right panel) hypotheses, β should be 0.5 and 1, respectively. The ellipse is the 0.95 quantile of covariance ellipse associated with the covariance matrix (upper right corner of the plot); the orange line is the scaling exponent fit through the constituent trait means.

To see why, note that a leaf’s *g*_max_ is determined by stomatal anatomy:

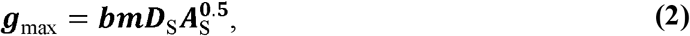

where *b* and *m* are biophysical and morphological constants, respectively (22) (see Methods for equations to calculate these constants). By Eq. 2, a higher *g*_max_ can be achieved with a smaller total stomatal area by increasing stomatal number and reducing stomatal size, because smaller stomata also have a shorter channel for diffusion. For example, consider two leaves with stomatal densities 250 and 200 pores mm^-2^ and stomatal areas 150 μm^2^ and 187.5 μm^2^. They have identical *f*_S_, but *g*_max_ at 25 °C is 11% greater for the leaf with smaller stomata (1.32 versus 1.47 mol m^-2^ s^-1^). Thus, selection for higher *g*_max_ would result in more numerous, smaller stomata, to minimize epidermal allocation to stomata and the evolution of higher *g*_max_ is strongly associated with the negative *A*_s_ and *D*_s_ relationship.

The ‘stomatal-area minimization’ hypothesis is controversial, however, as it is at odds with data in the literature that instead support an opposite, ‘stomatal-area increase’ hypothesis, i.e., that *g*_max_ should increase with *f*_S_ during evolution. Conversely, selection decreased *g*_max_ would be associated with decreased *f*_S_. The positive covariance of *g*_max_ with *f*_S_ has been shown in many studies and has been utilized in many papers in the literature that have indeed used *f*_S_ as a proxy for *g*_max_ (25, 26). According to the “stomatal-area increase” hypothesis, selection for higher *g*_max_ is much stronger than that to minimize cost, leading to greater surface allocation, even if this incurs a cost. Under this scenario, the negative *A*_s_ and *D*_s_ relationship would not act directly as a constraint on *g*_max_. Yet, selection for higher *g*_max_ would generate a distinctive covariation between these constituent anatomical traits. First, no relationship of *A*_s_ and *D*_s_ is absolutely required, which is consistent with data in the literature for species sets for which no relationship is found (27); as less than 50% of the leaf surface is typically taken up by stomata and many qualitatively different relationships between stomatal size and density across species are geometrically possible, including negative, zero, and positive covariances (ellipses in Fig. 1a). Yet, on average, a specific covariation would be expected if many combinations of *A*_s_ and *D*_s_ have similar fitness through their effect on either *g*_max_ or *f*_S,_ as we derive below.

It is critical to distinguish between these hypotheses for the evolution of *g*_max_ and the potential for *f*_S_ to constrain the observed stomatal size-density relationship. Implications of stomatal-area minimization are that *g*_max_ is ultimately constrained by the costs of high *f*_S_, that such costs are minimized, and that evolving higher *f*_S_ would be slowed by costs associated with allocating too much epidermal area to stomata^27–32^. By contrast, the stomatal-area increase hypothesis implies that selection on stomatal size and density primarily optimizes *g*_max_, which varies across environments, and greater *g*_max_ incurs stomatal construction costs and opportunity costs of epidermal space. Testing these hypotheses will further reveal how the evolution of high *g*_max_ relates to the general inverse stomatal size-density relationship.

Indeed, these hypotheses can be tested against data for diverse species by considering in detail the covariation among *D*_S_ and *A*_S_, for which they make different predictions. Under both hypotheses, *D*_S_ and *A*_S_ are constituents of composite traits, *f*_S_ or *g*_max_ (Eq. 1-2; Fig. 1b). We investigated how stomatal size-density scaling would differ between the hypotheses using models of macroevolutionary landscapes (28-31). We used the Ornstein-Uhlenbeck (OU) model originally derived from quantitative genetics for intraspecific (population) trait microevolution by Lande (32), and developed by Hansen (29)and others (28) for macroevolutionary interspecific trait variation. In the macroevolutionary OU model, interspecific trait variation expands through time until it reaches a stationary distribution around a long-term average(29). Within each species, microevolutionary forces (selection, genetic drift, mutation, and migration) and the environment drive genetic and plastic trait variation, respectively, and species’ trait means should be near their current adaptive optimum. The across-species distribution that becomes stationary in the OU model is thus dependent on these independent shifts in species’ optimum trait values. At stationarity, an OU process leads to stable trait mean and variance, setting the overall phenotypic constraint. Fitness tradeoffs likely limit the breadth of values for adaptive trait optima, given that extreme trait values will rarely optimize competing functions (33). Notably, the specific mechanisms for constraints on trait values are not specified but are implicit in the application of Ornstein-Uhlenbeck (OU) process to model evolution phenomenologically.

Given that the stomatal-area minimization and increase hypotheses differ in their prediction of how the species variation in composite traits (*f*_S_ and *g*_max_) are constrained by their constituent traits (*D*_S_ and *A*_S_), examination of the trait evolution can indicate which hypothesis was supported. The OU model can indicate *which* composite trait, *f*_S_ or *g*_max_, is primarily constrained. In both cases, analogous quantitative theory shows that constraint on composite traits imposed by stabilizing selection limits variation in constituent traits(34), and constraint on *f*_S_ results in a different covariance structure of *D*_S_ versus *A*_S_ than a primary constraint on *g*_max_. Note that both *f*_S_ and *g*_max_ show similar mathematical dependence on *D*_S_ and *A*_S_:

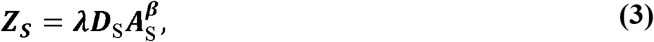

where composite stomatal trait *Z*_S_ (i.e., *f*_S_ or *g*_max_) is proportional to the product of constituent stomatal traits, with scaling exponent *β* multiplied by a scalar *λ*, which reflects stomatal dimension proportionalities and physical diffusion factors (22). For *g*_max_, *λ* = *bm* and *β* = 0.5 (Eq. 1); for *f*_S_, *λ* = 1 and *β* = 1 (Eq. 2). Since all traits are log-normally distributed^31^, and the OU model assumes Gaussian traits, we log-transformed Eq. 3:

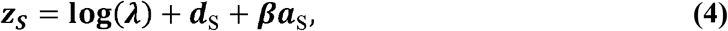

where lowercase variables indicate log-transformation of uppercase counterparts. Log-transformation also has the advantage of simplifying variance decomposition by linearizing the equation and enables traits measured on different scales to be directly compared in their proportional changes. Using random variable algebra, the variance in *z*_S_ is defined as:

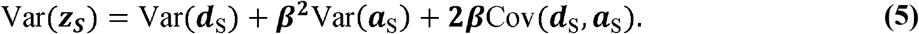

Using the variance-covariance of *d*_S_ and *a*_S_, we can find the scaling exponent *β* that minimizes Var(*z*_S_):

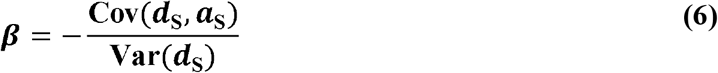

Notably, the right-hand side of Eq. 6 is the negative of the ordinary linear regression slope of log-stomatal size against log-density. Thus, for any dataset, *β* can be estimated using ordinary regression methods, but a negative slope estimate will result in a positive value of 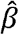. The stomatal-area minimization hypothesis predicts that 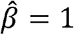 because *f*_S_ constrains *d*_S_ and *a*_S_ (Eq. 1), whereas the stomatal-area increase hypothesis predicts that 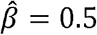 because *g*_max_ constrains *d*_S_ and *a*_S_ (Eq. 2). Note that the above prediction assumes that the primary constrained composite trait will also be the least variable composite trait, which allowed to identify the relationship between *β* and trait (co)variance in Eq. 6. We evaluated this assumption using forward-time, individual based, macroevolutionary quantitative genetic simulations (Supplementary Information). In each simulation, 1000 independent lineages evolve toward a moving optimal composite trait until stationarity following an OU process. The simulations confirm that the constrained composite trait is the least variable and that ordinary regression on interspecific trait means can accurately identify the simulated *β*. Estimates of *β* are not substantially affected by microevolutionary details about mutational and genetic covariances or geometric constraints on *f*_S_ (Fig. S2-S5).

We estimated stomatal size-density scaling in 2408 forest plant species from new field-collected samples over 28 sites in China and global synthesis of data from the literature (Fig. 2) and estimated the scaling exponent *β* using OU phylogenetic multiple regression with group (Angiosperm, Pteridophyte, Gymnosperm) and growth form (tree, shrub, herb) as covariates (see Methods).

**Fig. 2.**
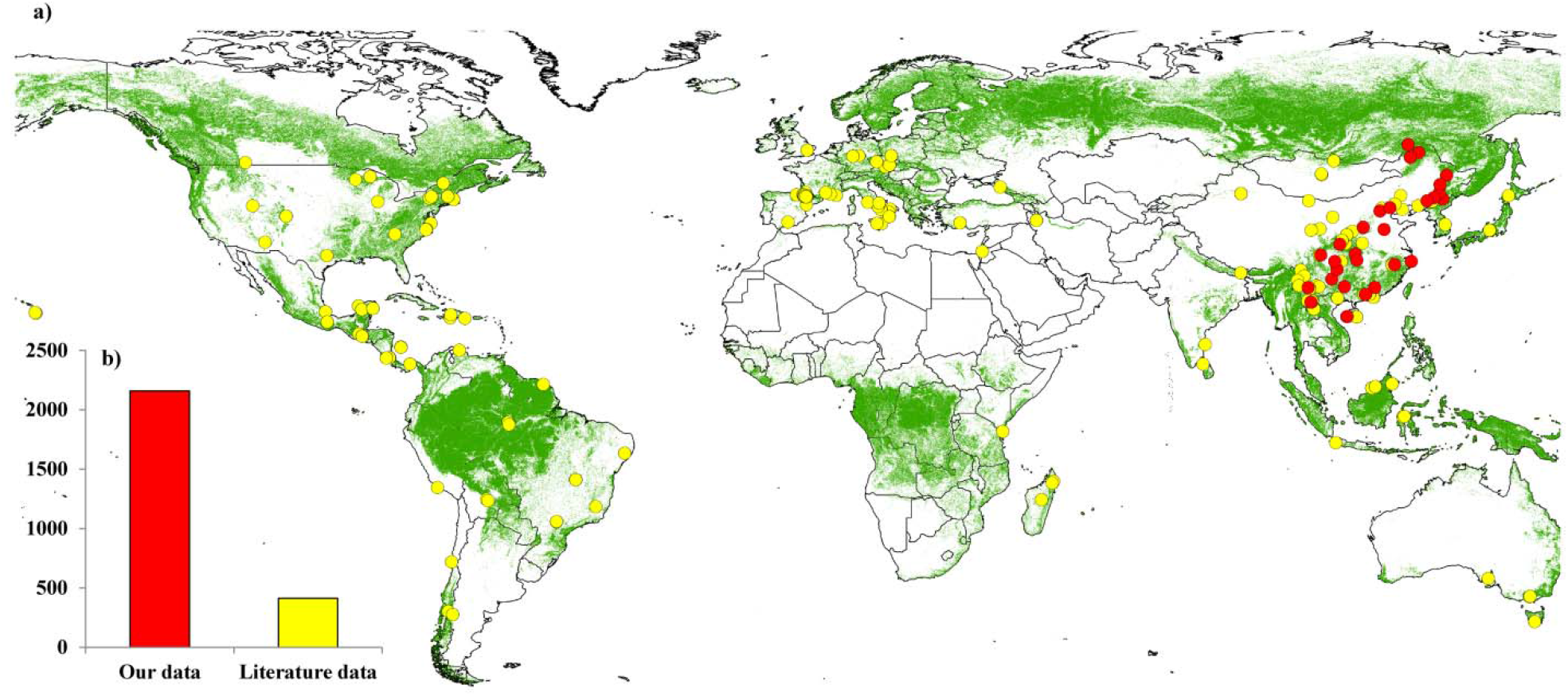
Geographic distribution of sampling sites (a) and the number of plant species (b) in this study.

Stomatal size-density scaling among forest plant species was consistent with a primary constraint on *g*_max_ (stomatal-area increase hypothesis, *β* = 0.5). Given the variance in stomatal density, the covariance between size and density among forest species minimizes the variance in *g*_max_. This implies that selection for higher *g*_max_ results in increased stomatal area allocation, and not minimizing area allocation (Fig. 3). There is no evidence that scaling differs between major groups, Angiosperms, Gymnosperms, and Pteridophytes (Fig. 3a; Table S1), but *g*_max_ is 49% (17-88% 95% CI; *P* = 0.001) and 14% (1-30% 95% CI; *P* = 0.04) higher in Angiosperms than Gymnosperms and Pteridophytes, respectively (Table S2). Trees also have 18% (8-28% 95% CI; *P* < 0.0001) and 48% (39-59% 95% CI; *P* < 0.0001) greater *g*_max_ than shrubs and herbs, respectively (Table S2). The across-species mean and variance in log(*g*_max_) are nearly invariant across latitude, temperature, and precipitation gradients, indicating that most of the variation in *g*_max_ occurs for species of contrasting ecology within rather than between forest sites, a finding similar to that for other key functional traits such as leaf mass per area and wood density (35) (Fig. 4).

**Fig. 3.**
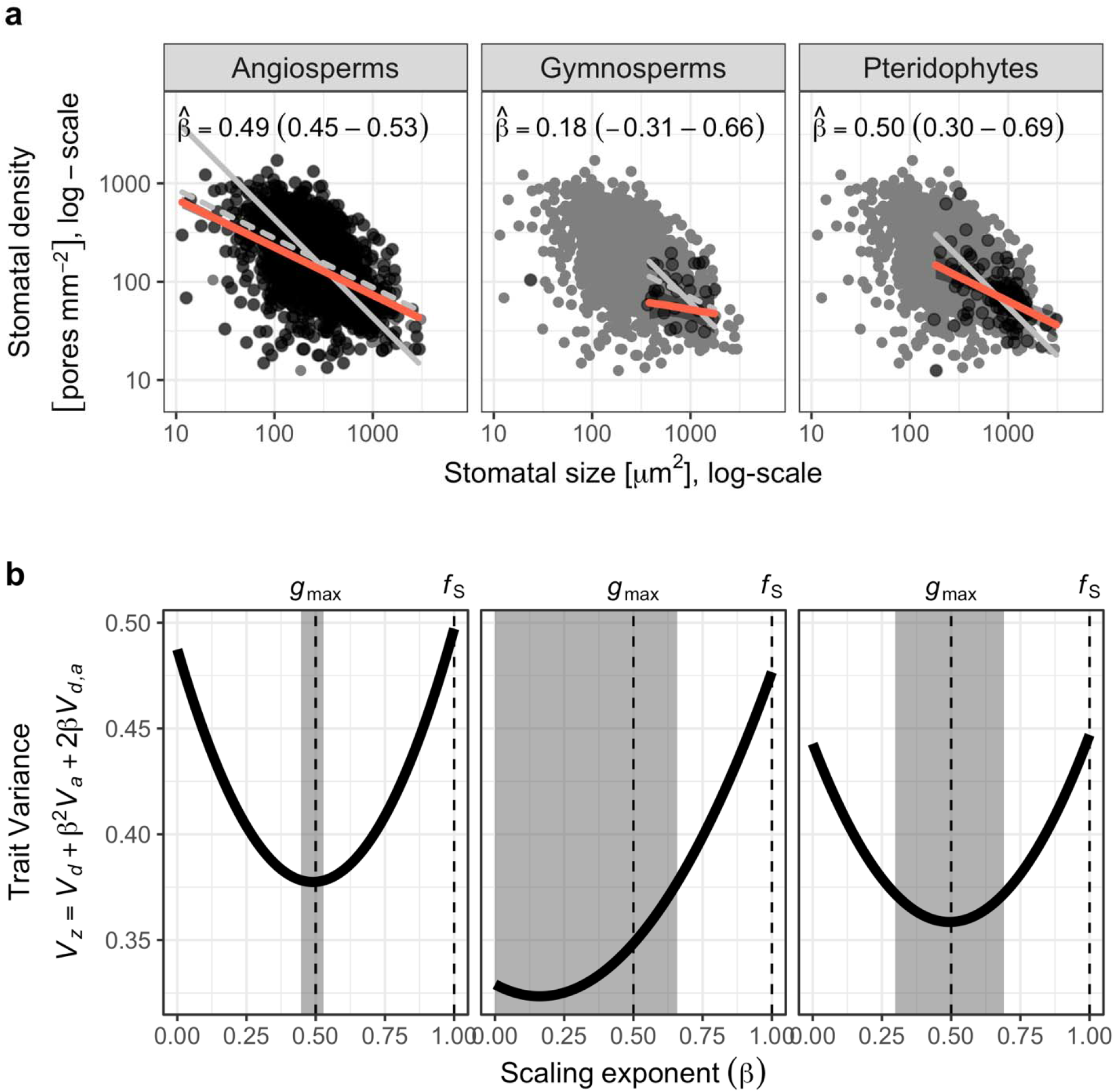
Stomatal size-density scaling is consistent with stomata-area increase but not area-minimization. **a**. In both angiosperms (left panel) and pteridophytes (right panel), the scaling exponent () estimated as the phylogenetic linear regression slope of stomatal size against density (Methods) is close to 0.5 as predicted by the stomatal-area increase hypothesis, but much less than 1.0, as predicted by the stomatal-area minimization hypothesis. For comparison, thin gray lines in the background show predicted slopes for each group when = 1.0 (solid line) and *β* = 0.5 (dashed line). The bootstrap 95% confidence intervals are in parentheses and shown graphically by the width of the grey rectangle in **b**. Dark points represent species mean trait values from the focal group; grey background points are from all groups for comparison. Orange line and ribbon are the estimated phylogenetic regression line and the 95% bootstrap confidence intervals. Scaling in gymnosperms (middle panel) is not significantly different from 0 or 0.5, but the confidence intervals do not include 1.0. **b**. The variance of the composite trait (*V*_*z*_) is minimized near β = 0.5, as predicted under the stomatal-area increase hypothesis (dashed-line under *g*_max_) but not where β = 1.0 as predicted by the stomatal-area minimization hypothesis (dashed-line under *f*_S_).

**Fig. 4.**
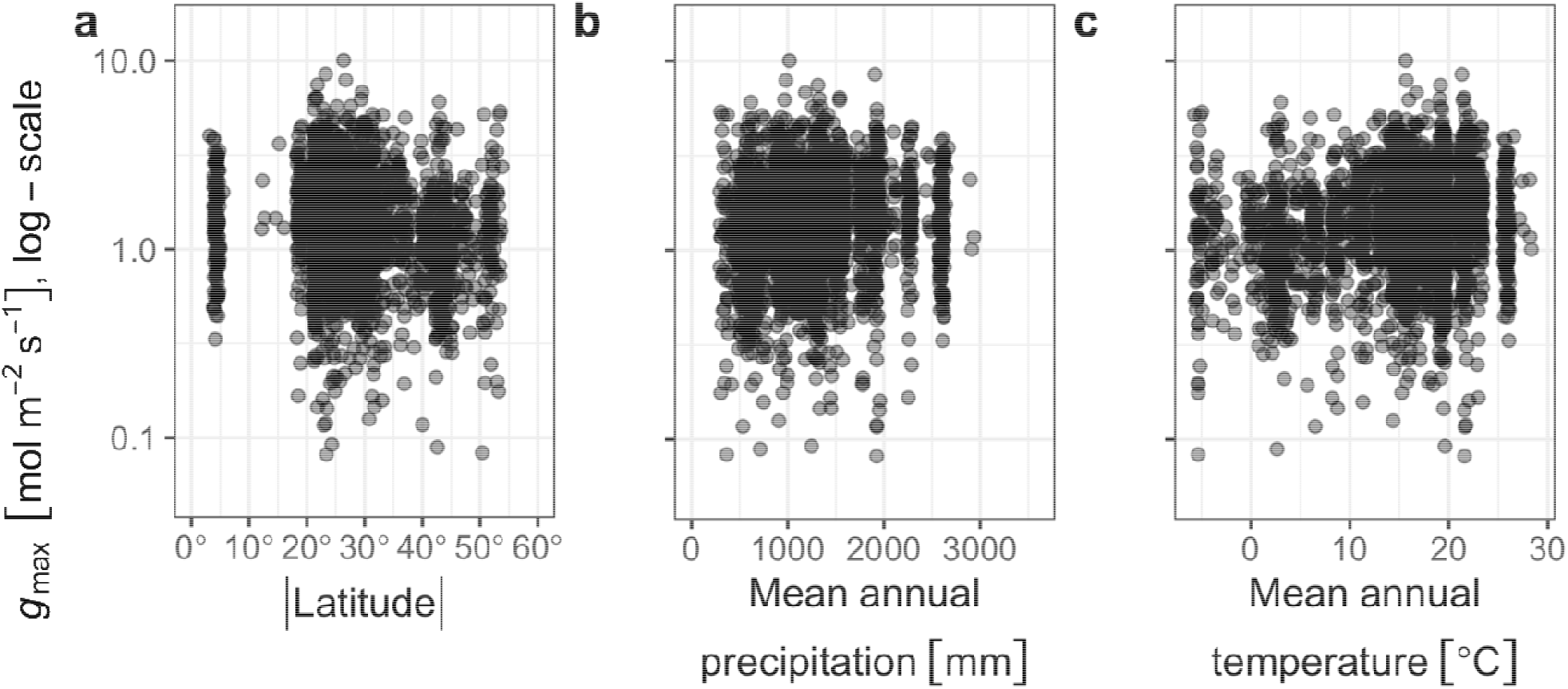
Anatomical maximum stomatal conductance varies little with latitude, mean annual precipitation, or mean annual temperature. Each point is the species’ mean |latitude| (**a**.), mean annual precipitation (**b**.), or mean annual temperature (**c**.) on the *x*-axis and the maximum stomatal conductance (*g*_max_) on the *y*-axis (log-scale). Based on phylogenetic multiple regression, the relationship between log(*g*_max_) and mean |latitude| (*P* = 0.69) and mean annual temperature (*P* = 0.10) are not significant; the relationship with mean annual precipitation is significant (*P* = 0.009) but weak since the total model *R*^2^ including all climate, lineage, and growth explanatory variables is only 0.11.

Our results overturn the prevailing view that the evolution of high *g*_max_ across diverse species is constrained by size-density scaling and minimized stomatal area allocation. Instead, the covariance between stomatal size and density supports stomatal area allocation increasing with the evolution of high *g*_max_. Thus, limits on the fraction of epidermis allocated to stomatal (*f*_S_) are a secondary consequence of limits on *g*_max_. Our novel analysis developed from quantitative genetic and macroevolutionary theory could distinguish the *g*_max_ evolution hypotheses. Notably, our *β* exponent for the scaling of *d*_S_ and *a*_S_ depends on using (phylogenetic) least squares regression, and thus, the results of studies reporting stomatal scaling slopes using standardized major axis (SMA) regression (which minimizes residual variance in both *d*_S_ and *a*_S_) would need to be recalculated to test against our findings (see Supplementary Information). Although estimated scaling using standard phylogenetic regression approaches (see Methods), it is more appropriate to interpret our results not as minimizing residual variance, but rather estimating the *β* consistent with the covariance structure of stomatal size and density (Fig. 1).

Our results have at least two important implications for understanding the evolutionary anatomical mechanisms of high *g*_max_ and its consequences for the stomatal size-density scaling relationship. First, the finding that size-density scaling does not constrain the evolution of higher *g*_max_ implies that stomatal cost is not a constraint on high *g*_max_ and thus a different constraint on the evolution of extreme values of *g*_max_ across environments. Very high *g*_max_ may be rare because the *g*_op_:*g*_max_ ratio is constrained in a region of maximal control to respond rapidly to changing environments (36). Additionally or alternatively, a high *g*_max_ may also be linked with a high wilting point thereby setting a physical upper limit to leaf gas exchange and a high risk of hydraulic failure (37) if open stomata face transiently high atmospheric drought. Other possible costs include detrimental consequences of high *g*_max_ for stomatal movements and diffusion, as well as energetic costs of opening closing more and/or larger stomata (38, 39). Future work should prioritize identifying the fitness costs and functional trade-offs that constrain the evolution of high *g*_max_. Second, if *g*_max_ is the primary constraint, this implies that space allocation to stomata is relatively unimportant, such that plants could allocate a greater fraction of their epidermal area to stomata than they currently do without counterveiling selection. Thus, if stomatal size and density can be manipulated independently, anatomies with the same *g*_max_, but different *f*_S_, would have similar fitness in the same environment. This finding also clarifies the evolution of stomata across major plant lineages, and refutes the hypothesis that smaller stomata were required to increase *g*_max_ in angiosperms (20). All three major land plant lineages have similar variance in *g*_max_ (Fig. 3b); angiosperms have higher *g*_max_ than gymnosperms and pteridophytes on average due to their higher *d*_S_ for a given *a*_S_, not because of differences in the scaling relationship. The higher stomatal density of angiosperms would be linked to increases in leaf water transport capacity, for example, by decreasing the distance between vein and stomata, allowing stomata to stay open^40^. The primary constraint on maximum stomatal conductance appears to be that selection rarely favors extreme values, implying that vegetation and crop models should incorporate nonepidermal costs of extreme trait values to predict optimal *g*_max_ for the prediction of photosynthetic carbon gain and transpiratory water loss across scales.

## Methods

### Stomatal trait data from global forests

The stomatal dataset of global forests represents a total of 2408 plant species from natural forests, including novel field data collected from Chinese forest communities and a compilation of published literature values.

Our field data were collected from 28 typical forest communities occurring between 18.7 °N and 53.3 °N latitude in China. The field sites were selected to cover most of the forest types in the northern hemisphere, including cold-temperate coniferous forest, temperate deciduous forest, subtropical evergreen forest, and tropical rain forest (Fig. 2). In total, we sampled 28 forest sites. We used the Worldclim database (40) to extract additional data on mean annual temperature (MAT) and precipitation (MAP) over the period 1960-1990 using latitude and longitude. Among these forests, mean annual temperature (MAT) ranged from -5.5-23.2 °C, and mean annual precipitation (MAP) varied from 320 to 2266 mm. The field investigation was conducted in July-August, during the peak period of growth for forests. Sampling plots were located within well-protected national nature reserves or long-term monitoring plots of field ecological stations, with relatively continuous vegetation. Four experimental plots (30 × 40 m) were established in each forest.

Leaves from trees, shrubs, and herbs were collected within and around each plot. For trees, mature leaves were collected from the top of the canopy in four healthy trees and mixed as a composite sample. Eight to 10 leaves from the pooled samples were cut into roughly 1.0 × 0.5 cm pieces along the main vein, and were fixed in formalin-aceto-alcohol (FAA) solution (5 ml 38 % formalin, 90 ml 75 % ethanol, 5 ml 100 % glacial acetic acid, and 5 ml 37 % methanol) (41). In the laboratory, three small pieces were randomly sampled, and each replicate was photographed twice using a scanning electron microscopy (Hitachi SN-3400, Hitachi, Tokyo, Japan) on the lower surface at different positions. We focused on the lower epidermis (42), because a previous study has demonstrated that most of leaf upper epidermis has no stomata for forest plants (43).

In each photograph, the number of stomata was recorded, and *D*_S_ was calculated as the number of stomata per unit leaf area. Simultaneously, five typical stomata were selected to measure stomatal size using an electronic image analysis equipment (MIPS software, Optical Instrument Co. Ltd., Chongqing, China).

Peer-reviewed papers on leaf stomata were collected using an all-databases search of Web of Science (www.webofknowledge.com) from 1900 to 2018 using “forest” and “stomata” as a topic, in line with the principle of “natural forest, non-intervention, species name” (i.e. we did not use data from controlled experiments or where taxonomic data was missing). A total of 90 papers (see Supporting Table S3) which met our requirements, yielding D_S_ and *L* measurements from 413 plant species (Fig. 2) from which we calculated *g*_max_ and *f*_S_. *f*_S_ is proportional to the stomatal pore area index (SPI), which defined as the product of *D*_S_ and stomatal length (*L*) squared (25), because *A*_S_ = *mL*^2^ (22).

We calculated *g*_max_ (Equation 1) to water vapor at a reference leaf temperature (T_leaf_ = 25° C) following Sack and Buckley (22). They defined a biophysical and morphological constant as:

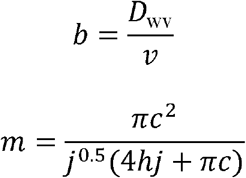

*b* is the diffusion coefficient of water vapor in air (*D*_wv_) divided by the kinematic viscosity of dry air (*v*). *D*_wv_ = 2.49 × 10^−5^ m^2^ s^−1^ and *ν* = 2.24 × 10^−2^ m^3^ mol^−1^ at 25° (44). For kidney-shaped guard cells, *c* = *h* = *j* = 0.5; for dumbbell-shaped guard cells in the Poaceace, *c* = *h* = 0.5 and *j* = 0.125. We used the species average *g*_max_ and *f*_S_ for all analyses.

### Phylogenetic regression

By positing that the least variable composite of stomatal size and density indicates the trait with the most constraint (Fig. 1), we identify a new way to estimate the scaling exponent *β* (Eq. 6) using linear regression estimates, and also accounted for phylogenetic nonindependence. We used the Plant List (http://www.theplantlist.org) to confirm species names, then we assembled a synthetic phylogeny using S.PhyloMaker (45). We fitted phylogenetic regression models using the **phylolm** version 2.6 package in R (46). As we derived in the main text, the scaling exponent *β* can be estimated from the slope of the regression of *a*_S_ on *d*_S_, where 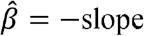. We estimated separate scaling exponents for major groups, Angiosperms, Pteridophytes, and Gymnosperms. We also estimated different intercepts, corresponding with different average *g*_max_ values, for functional types (herbs, shrubs, and trees) and grasses, because of their unique stomatal anatomy. We used the “OUrandomRoot” model of trait evolution. 95% confidence intervals for all parameters were estimated from 1000 parametric bootstrap samples generated by simulating from the best-fit model and re-fitting. *P*-values for coefficients are based on *t*-tests. We used the same methods to test whether *g*_max_ (log-transformed for homoskedasticity) was affected by |latitude|, MAP, MAT, group (Angiosperms, Pteridophytes, Gymnosperms), and/or functional type (herb, shrub, tree). One gymnosperm species, *Torreya fargesii*, had substantially lower stomatal size than would be predicted from its density (Fig. 3a). There results of the paper did not change if this outlier was excluded because the confidence intervals for stomatal-density scaling are very wide for Gymnosperms regardless. Therefore, we excluded this species from statistical analyses but show it in the figure for completeness. All data were analyzed in R (47)version 4.0.5

## Acknowledgements

Financial support was supported by the National Natural Science Foundation of China (31988102, 31770655, 31870437), the National Key R&D Program of China (2017YFA0604803), the second Tibetan Plateau Scientific Expedition and Research Program (2019QZK060202), The Chinese Academy of Sciences Strategic Priority Research Program (XDA23080401), US National Science Foundation 1929167 (to CDM), and the Project funded by China Postdoctoral Science Foundation (2020M680663). We thank “Functional Trait Database of Terrestrial Ecosystems in China (China_Trait)” for sharing data, further information for other materials should contact to N.P. He. There are no conflicts of interest to declare.

## Author contributions

N.H. and G.Y. designed field sampling; N.H., C.D.M, and C.L. conceived the initial ideas; C.L., N.H., Y.L. J.Z., Z.Z., M.L. and L.X collected the data; C.L. wrote the first draft, and C.D.M. contributed the final mathematical derivations, data analysis, and wrote the final manuscript; L.S., H.J.B., C.L., N.H., G.Y., and X.H. revised the manuscript. All authors gave final approval for publication.

## Supplementary information

