## Supplemental Material 1 for "Scaling between stomatal size and density in forest plants" (from another version of this preprint: version 1, 2021-04-26)

**Supplementary Information**

*Estimating stomatal size-density scaling*

Standard approaches to model and estimate scaling exponents borrowed from allometry^1^ are not the correct framework for distinguishing hypotheses for stomatal size-density scaling. Allometry traditionally compares the relative growth of two biological entities (e.g. brain versus body mass or root versus leaf mass). Researchers predict scaling exponents (*β*) relating one size (*M*_1_) to another (*M*_2_) using optimality or geometric models:

$$\boldsymbol{M}_{\boldsymbol{2}}\boldsymbol{=A}\boldsymbol{M}_{\boldsymbol{1}}^{\boldsymbol{\beta}}$$

In contrast, the hypotheses for stomatal size-density scaling posit that constraint on a composite trait ($Z_{S}={\lambda D}_{\text{S}}A_{\text{S}}^{\beta}$; Eq. 3) shapes the covariance of constituent traits (*D*_S_ and *A*_S_). Despite the mathematical similarity of allometric and size-density scaling, the hypotheses to explain these relationships are subtly, but fundamentally different. In allometry, hypotheses usually posit that selection on scaling or a mechanistic link constrains the relationship between *M*_1_ and *M*_2_; in stomatal scaling, constraints on the variance of a composite trait *Z*_S_ generate a scaling relationship between *D*_S_ and *A*_S_ as a byproduct.

These fundamentally different hypotheses for scaling relationships render the arguments for using standardized major axis (SMA) regression in allometric studies unsuited estimate constraint on stomatal size-density scaling. To illustrate this point, we linearize the allometric equation using log-transformation:

$$\boldsymbol{m}_{\boldsymbol{2}}\boldsymbol{=a+\beta}\boldsymbol{m}_{\boldsymbol{1}}$$

For brevity, lowercase variables are the log-transformed counterparts of uppercase variables. Allometric studies usually want to test whether a “line-of-best-fit” ^1^ is consistent with a single predicted *β* because models often predict a single optimal *β* to explain how *m*_1_ and *m*_2_ covary. Variance away from the optimal relationship are attributed to variance in the intercept (*a*) or variance in the scaling exponent *β*. We ignore measurement error here because we are concerned with broad comparative studies in which measurement error is small compared to biological differences. Estimating the “line-of-best-fit” depends on assumptions one makes about which residual variance to minimize. Linear and SMA both fit the data equally well, but estimate slopes differently. Linear regression minimizes residual variance in the response variable; SMA minimizes residual variance both explanatory and response variables, scaled. SMA is usually favored in allometry studies because *m*_1_ and *m*_2_ do not bear a cause-and-effect relationship, but SMA slopes may be biased or unstable when the variance explained is low^2^. Both linear and SMA regression can incorporate phylogenetic information.

The distinguishing feature of stomata-area minimization and increase hypotheses for stomatal size-density scaling is not how well lines with different slopes fit the data. Rather, as described in the main text, we posit that the composite trait *Z*_S_ with the least variance is the most constrained because of its functional significance. As we show, the (phylogenetic) linear regression slope is also an estimate of the *β* that minimizes *Z*_S_, but this should not be interpreted as a “line-of-best-fit” because that is not the distinguishing feature of these competing hypotheses. There are multiple lines that fit the data equally well, depending on which residuals are minimized, but different slopes.

Because the linear regression slope is an estimator of *β*, the SMA slope used in allometric studies is biased. For example, de Boer *et al.* (2016) estimated a size-density scaling slope of ~1 using SMA approaches based on allometry studies^3^. The linear regression slope estimate is simply the SMA slope multiplied by the sample correlation coefficient (*r_xy_*):

$$\text{SMA slope = ±}\frac{s_{y}}{s_{x}}$$

$$\text{Linear regression slope = }r_{xy}\frac{s_{y}}{s_{x}}$$

where *s_x_* and *s*_y_ are the sample standard deviations of *x* and *y*, respectively. These methods will estimate scaling exponents the same when the correlation is perfect (|*r_xy_*| *=* 1); otherwise SMA slope will be more biased as |*r_xy_*| goes to 0. More importantly, the SMA slope is not sensitive to the covariance whereas the regression slope is, through the correlation coefficient. This is important because the competing biological hypotheses for stomatal size-density scaling specifically posit that constraint on the composite trait shapes covariance in the constituent traits.

To illustrate how SMA regression biases estimates of *β*, we calculated the expected result over a range of plausible parameters. We assume that the evolutionary variance of the composite stomatal trait (log-scaled) *z*_S_ is unity (Var(*z*_S_) = 1), so that other trait variances can be interpreted as relative to variance in *z*_S_. For example, if the variance of log-density (*d*_S_) is 0.5, this scenario can be rescaled to any situation where the variance of *d*_S_ is half that of *z*_S_. The variance of *z*_S_ represent the range of adaptive optima among extant woodland plants, assuming most populations are near their current adaptive optima. The range of adaptive optima is constrained by limited area (stomatal-area minimization hypothesis, *β* = 1), selection against extreme *g*_max_ (stomatal-area increase, *β* = 0.5), or another factor or factors (*β* ≠ 0.5 or 1).

In the calculations below, we set Var(*z*_S_) to unity (see above) and the relative variance in *a*_S_ varied between 0.5 and 2. Once Var(*z*_S_) and Var(*a*_S_) are specified, we can calculate the expected Var(*d*_S_) and Cov(*d*_S_, *a*_S_) for a given *β* (0.5 or 1) following Eq. 4, assuming that $\beta=-\frac{\text{Cov}(d_{\text{S}}, a_{\text{S}})}{{\text{Var}(d}_{\text{S}})}$ (see main text). The resulting (co)variance relationship ensures that the composite trait variance with the given *β* will have lower variance than all other composite traits derived with another *β* value. Note that we could have equivalently specified Var(*d*_S_) then calculated Var(*a*_S_) and Cov(*d*_S_, *a*_S_)

Next, we show that the linear regression (LR) slope, by definition, accurately estimates the scaling relationship under each hypothesis. To reiterate the main text, estimated *β* log-stomatal size against log-density times -1. In contrast, the SMA slope (see above, also multiplied by -1 to estimate *β*) and Pearson correlation coefficient vary considerably depending on the variance in constituent traits (*a*_S_ and *d*_S_). Note that we use phylogenetic (co)variances in the analysis to estimate regression slopes in the main text, which is conceptually the same as LR except they account for phylogenetic nonindependence. We could calculate phylogenetic SMA slopes using the same adjustment. For simplicity, we demonstrate the difference between nonphylogenetic LR and SMA here, but the exact same relationships apply to phylogenetic estimates. Over reasonable parameter ranges, SMA slopes also significantly overestimate *β*, whereas the LR slope (by definition) is an unbiased estimator of -*β*. We compared these theoretical calculations with the actual data by plotting the variance of *a*_S_, *d*_S_, their covariance relative to the variance in *g*_max_ and *f*_S_. These calculations are approximate because we did not factor out growth form or account for phylogenetic nonindependence, as we did in the main analysis. Nevertheless, they illustrate that the covariance structure is much closer to that predicted by stomatal-area increase hypothesis (Fig. S1).

**Tables**

**Table S1 | Stomatal size-density scaling is preserved across major plant lineages.** In this phylogenetic regression model we looked at the slope between stomatal size (log-scale) and density (log-scale) across Angiosperms, Gymnosperms, and Pteridophytes. We also estimated separate intercepts for each group as well as grasses, shrubs, and trees. Estimates are derived from phylogenetic least squares estimation; 95% confidence intervals are based on 1000 parametric bootstrap simulations. Ornstein-Uhlenbeck (OU) phylogenetic regression parameters model the residuals. We list the individual slopes for each group, as these are estimates of the scaling exponent. The overall intercept for Angiosperm herbs, so the coefficients for phylogenetic groups and functional types effect their intercept relative to Angiosperms herbs.

| **Term** | **Estimate** | **95% Confidence interval** |
| --- | --- | --- |
| Slope $(=-\hat{\beta})$: |  |  |
| Angiosperms | -0.49 | -0.53 – -0.45 |
| Gymnosperms | -0.18 | -0.66 – 0.31 |
| Pteridophytes | -0.50 | -0.69 – -0.30 |
| Intercept  (Angiosperm herb): | 7.65 | 7.42 – 7.89 |
| Effect of Gymnosperm | -2.59 | -5.75 – 0.78 |
| Effect of Pteridophyte | -0.10 | -1.45 – 1.27 |
| Effect of Grass | -0.87 | -1.05 – -0.69 |
| Effect of Shrub | 0.25 | 0.18 – 0.31 |
| Effect of Tree | 0.41 | 0.34 – 0.47 |
| OU Parameters: |  |  |
| α | 1.56 | 1.01 – 2.73 |
| σ^2^ | 1.17 | 0.75 – 2.04 |

**Table S2 | Anatomical maximum stomatal conductance (*g*_max_) is similar across latitude and climate.** Latitude (absolute value), mean annual precipitation (MAP) and mean annual (MAT) of species had no significant effect on *g*_max_ (log-scale). Here we report the estimated slope and 95% confidence interval for each of these continuous explanatory variables regressed against *g*_max_ (log-scale). Gymnosperms and Pteridophytes have lower *g*_max_ than angiosperms in the same functional group. Shrubs and trees have higher *g*_max_ than herbs within the same group. Estimates are derived from phylogenetic least squares estimation; 95% confidence intervals are based 1000 parametric bootstrap simulation. OU parameters model the phylogenetic terms of the residuals.

| **Term** | **Estimate** | **95% Confidence interval** |
| --- | --- | --- |
| Intercept (Angiosperm herb) | 0.17 | 0.31 – 0.62 |
| \|latitude\| | -1.87 × 10^-3^ | -10.7 – 7.50 × 10^-3^ |
| MAP | -1.19 × 10^-4^ | -1.99 – 0.00 × 10^-4^ |
| MAT | 8.59 × 10^-3^ | -1.95 – 18.6 × 10^-3^ |
| Gymnosperms | -0.40 | -0.62 – -0.16 |
| Pteridophytes | -0.13 | -0.26 – -0.006 |
| Grasses | -0.55 | -0.72 – -0.36 |
| Shrubs | 0.23 | 0.17 – 0.30 |
| Trees | 0.40 | 0.34 – 0.46 |
| OU Parameters: |  |  |
| α | 1.44 | 0.88 – 2.95 |
| σ^2^ | 1.92 | 1.16 – 3.91 |

**Figures**

**Fig. S1 | Standardized major axis (SMA) regression slopes overestimate stomatal size-density scaling.** In these calculations, we assume that Var(*z*_S_) = 1, so that the (co)variance of constituent traits (Var(*a*_S_), Var(*d*_S_), and Cov(*d*_S_*,a*_S_)) are *relative* to the variance in *z*_S_. See ‘Estimating stomatal size-density scaling” above. **a.** We calculated the correlation (red, Cor(*d*_S_*,a*_S_)) covariance (green, Cov(*d*_S_*,a*_S_)), and Var(*d*_S_) (blue), for *β* = 0.5 (left, stomatal-area increase) and *β* = 1 (right, stomatal-area minimization). The estimated values (points) are closer to expectations for stomatal-area increase, consistent with our estimates that *β* = 0.5. **b.** This shows that linear regression slopes are an unbiased estimate of *β*, whereas SMA slopes overestimate *β*.
