## Supplemental Material 2 for "Scaling between stomatal size and density in forest plants" (from another version of this preprint: version 1, 2021-04-26)

### Supplementary Information: Macroevolutionary quantitative genetic simulations

To reiterate, we hypothesized in the main text that selection acts primarily on a composite trait ($f_S$ or $g_{\text{max}}$). Scaling between constituent traits (stomatal density and size) is a byproduct of constraint on the composite trait. Our derivation implicitly assumes no other factors constrain the (co)variance of constituent traits, which is a strong simplification. Here we consider three factors which affect the (co)variance of constituent traits and could therefore alter our predictions: genetic drift, mutation, and physical limits. Genetic drift decreases trait variance and can cause populations to evolve away from their adaptive optimum. Mutation increases trait variance and can alter trait covariance if mutations have pleiotropic effects on multiple traits. Finally, physical limits reduce trait variance. For example, stomatal cover by definition cannot exceed 1. Here we complement our analytic theory using forward-time, individual-based, quantitative genetic simulations to evaluate whether these additional factors (drift, mutation, and physical limits) alter our primary inference that $\beta$ can be estimated from the scaling between constituent traits across species. Note that our purpose is not to realistically simulate the evolution of these traits in land plants or use simulation results as a basis for estimating quantitative genetic parameters from phylogenetic comparative data.

#### Macroevolutionary quantitative genetic model

In this section we describe a macroevolutionary quantitative genetic modeling framework; in the next section we describe simulations we used to evaluate the effects of drift, mutation, and physical limits on scaling between constituent traits. The model integrates macroevolutionary (between species) and microevolutionary (within species) approaches. Between species we use macroevolutionary theory to simulate how the adaptive landscape changes through time; within species, we use quantitative genetic theory to simulate the evolution of continuous phenotypes using phenomenological variance components. Below, we describe how

1. traits are determined by genetic and environmental contributions,
2. the macroevolutionary adaptive landscape changes through time, and
3. selection, drift, and mutation determine microevolution within species.

#### Traits are determined by genetic and environmental contributions

We model a composite trait $z$ and its constituent traits $x$ and $y$ as random variables with different constants, $z_0$ and $\beta$:

$$z = z_0 + x + \beta y \tag{1}$$

This equation is structurally identical to Eq. 4 in the main text if $x$ is the log of stomatal density ($d_\mathrm{S}$) and $y$ is the log stomatal size ($a_\mathrm{S}$). $z$ is equivalent to $f_S$ when $z_0 = 0$ and $\beta = 1$; $z$ is equivalent to $g_\mathrm{max}$ when $z_0 = \log(bm)$ and $\beta = 0.5$.

Consider $P$ independent "species" each with a constant, identical effective population size $N$ undergoing mutation, selection, and genetic drift. Each species is panmictic and there is no gene flow between species, so they evolve as independent lineages from the start of time. Multivariate quantitative genetic models that combine deterministic (mutation and selection) and stochastic (drift) forces are generally not analytically tractable[1], therefore we use forward-time, individual-based, discrete-generation simulations. Hence, simulations track the composite trait value of individual $i$ in species $j$ at time $t$ as:

$$z_{ijt} = z_0 + x_{ijt} + \beta y_{ijt} \tag{2}$$

Constituent traits ($x_{ijt}$, $y_{ijt}$) are a function of additive genetic ($g$) and environmental variance ($e$):

$$\begin{aligned} x_{ijt} &= g_{x,ijt} + e_{x,ijt} \\ y_{ijt} &= g_{y,ijt} + e_{y,ijt} \end{aligned} \tag{3}$$

For simplicity, we ignored nonadditive genetic variance components, which aren't heritable but can effect trait expression. $g_{x,ijt}$ and $g_{y,ijt}$ are drawn randomly from a multivariate normal distribution with mean vector $\boldsymbol{\mu}_{G,jt}$ and additive genetic covariance matrix $\mathbf{G}_{jt}$. We made no *a priori* assumptions about $\boldsymbol{\mu}_{G,jt}$ or $\mathbf{G}_{jt}$, but allowed them to evolve in each lineage through mutation, genetic drift, and selection. Environmental contributions $e_{x,ijt}$ and $e_{y,ijt}$ are drawn randomly from a multivariate normal distribution with mean vector $\vec{0}$ and environmental covariance matrix $\mathbf{E} = \mathrm{diag}(V_E)$. In other words, we assumed constant, uncorrelated environmental trait variance $V_E$. Since $\mathbf{E}$ is constant, trait heritability depends on the additive genetic variance within a lineage at a given time.

#### The macroevolutionary adaptive landscape changes through time

Within each lineage, the optimal composite trait value, $z_{\text{opt}}$, changes through time, independently in each lineage, following an Ornstein-Uhlenbeck (OU) process, which most adequately describes the dynamics of many trait optima[2].

$$z_{\text{opt}}[t+1] = \alpha(z_{\text{opt}}[t] - \theta) + \sigma W t \tag{4}$$

$\theta$ is the long-run average $z_{\text{opt}}$, $\sigma$ is the rate of random movement in the optimum each generation, and $\alpha$ is the rate of return to $\theta$. We assume that the OU process is homogenous through time in all lineages, meaning parameters are constant. The macroevolutionry OU process has been described extensively elsewhere[3–6]. Note that there are no independent adaptive optima for constituent traits. As we show below, the effect of constituent traits on fitness is completely mediated by their effect on the composite trait. For simulations, we discretized the OU process by randomly sampling $z_{\text{opt}}[t+1]$ using the `rOU` function from the *R* package **POUMM** version 2.1.7[7].

#### Selection, drift, and mutation determine microevolution within species

##### Selection

Each generation, species evolve toward the current optimal composite trait value $z_{\text{opt}}$. We assume a Gaussian fitness function:

$$W = W_{\max} e^{\frac{-(z - z_{\text{opt}})^2}{V_S}} \tag{5}$$

$W$ is absolute fitness, $W_{\max}$ is the maximum absolute fitness when $z = z_{\text{opt}}[t]$, and $V_S$ is the strength of stabilizing selection around $z_{\text{opt}}[t]$. For simplicity, we assume that $W_{\max}$ and $V_S$ are equal to 1 and are constant through time for all species. These parameters affect the rate of adaptive evolution within lineages and the composite trait variance, but should not affect the constituent trait covariance, which is our principal interest here.

##### Genetic drift

We introduce genetic drift by sampling individuals with replacement weighted by their relative fitness. The power of genetic drift to change the phenotype therefore depends on the population size relative to the genetic variance in fitness.

##### Mutation

Mutation occurs after selection and drift. Let $\mathbf{H}_{j,t}$ be an $N \times 2$ matrix of additive genetic contributions within species $j$ at time $t$ following selection and drift. The additive genetic contribution in the next generation is:

$$\mathbf{H}_{j,t+1} = \mathbf{H}_{j,t} + \mathbf{U} \tag{6}$$

where $\mathbf{U}$ is an $N \times 2$ matrix of mutational effects. Following standard quantitative genetic theory, we assume mutation is bivariate normal ($\mathcal{N}_2$) with a mean vector $\vec{0}$ and mutational covariance matrix $\mathbf{M}$:

$$\mathbf{U} \sim \mathcal{N}_2(\vec{0}, \mathbf{M}) \tag{7}$$

We assume that mutation is unbiased (mean vector is $\vec{0}$) and that $\mathbf{M}$ is constant through time in all species. $\mathbf{M}$ is a $2 \times 2$ covariance matrix with the diagonals equal to the mutational variance of constituent traits and the covariance on the off-diagonals:

$$\mathbf{M} = \begin{bmatrix} V_{M,X} & V_{M,XY} \\ V_{M,XY} & V_{M,YY} \end{bmatrix} \tag{8}$$

In simulations, we varied the mutational variances and correlation, $\rho_{M,XY} = V_{M,XY} / \sqrt{V_{M,X} V_{M,Y}}$.

#### Stationarity

We ran simulations until the between-species mean and trait covariance reached a stationary distribution, i.e. stationarity. The composite trait $z$ quickly reaches stationarity as a product of both random variation in $z_{\text{opt}}$ and within species variation around its $z_{\text{opt}}$. However, constituent traits $x$ and $y$ do not necessarily reach stationarity because there are no direct bounds on $x$ and $y$ except in the physical limits model (see below).

### Simulations

We conducted four sets of macroevolutionary quantitative genetic simulations to address how genetic drift, mutation, or physical limits change interspecific covariance between constituent traits (Table S3). The code to run all simulations will be made publically available upon publication at https://github.com/cdmuir/stomata-scaling.

| | Set 1 | Set 2 | Set 3 | Set 4 |
|---|---|---|---|---|
| Selection on composite trait | ✓ | ✓ | ✓ | ✓ |
| Genetic drift | ✓ | ✓ | ✓ | ✓ |
| Unequal mutational variance | | ✓ | | |
| Mutational correlation | | | ✓ | |
| Physical limits | | | | ✓ |

Table S3: Summary of simulation sets 1-4. All simulations include selection and genetic drift, but differ in how they model mutation and physical limits.

For each parameter combination in all sets, we simulated the evolution of $x, y,$ and $z$ in 1000 species until stationarity for cases when $\beta = 0.5$ and $\beta = 1.0$. We then estimated $\beta$ from simulated data following the same approach in our analysis. The only difference is that in our simulations, we used a star phylogeny in which all species diverge at the beginning of the simulation and are statistically independent, hence we used standard statistical approaches rather than phylogenetic approaches. If genetic drift, mutation, and physical limits do not affect our predictions, then we should be able to correctly estimate $\beta$ in simulated data sets.

#### Parameters

Table 2 summarizes all parameters used in each set of simulations. Here we discuss how we chose parameter values. In all sets, we simulated selection on composite traits where $\beta = 0.5$ and $\beta = 1$, the hypothesized scaling values. With the exception of Set 4, we arbitrarily set $z_0 = 0$ for simplicity.

In sets 1-3, we set macroevolutionary parameters so that the stationary distribution of $z$ would have mean $\theta = 0$ and variance $(\sigma^2/(2\alpha) = 0.5)$. For this set of simulations, the long-run average is arbitrary, but we set the variance close to the observed variance of $\log(g_{\max})$ and $\log(f_S)$, 0.42 and 0.54, respectively, to make it easier to compare results with Set 4 in which we use realistic trait values to apply physical limits. However, preliminary simulations did not find that the stationary variance had any effect on the ability to correctly estimate $\beta$ from simulated data (results not shown).

Preliminary analysis showed that a population size to $N = 1024$ was sufficiently large for populations to respond to selection and track the adaptive optimum with the macroevolutionary parameters discussed in the previous paragraph. Smaller $N$ increased the variability in simulations but had no qualitative effect; larger $N$ took more time without qualitatively changing the results (results not shown). Since we imposed soft selection where population size is constant regardless of absolute fitness, relative fitness determines the response to selection. Hence, we arbitrarily set the maximum absolute fitness $W_{\max} = 1$ in all simulations. In Sets 1-3, we arbitrarily set $V_E = 1$ and adjusted macroevolutionary parameters (see above) so that species could track the adaptive optimum through time. We explain differences with Set 4 below. Following standard quantitative genetic assumptions, we set $1000V_M = V_E$ and $V_S = 10V_E$ unless otherwise noted. These values correspond to weak mutation and moderate selection values that are realistic for most organisms[8–10]. We describe additional parameter choices that varied among simulations sets below.

##### Set 1: Selection and genetic drift

This is the baseline scenario with selection and genetic drift, as in the previous section, but equal genetic variances in constituent traits ($V_{M,X} = V_{M,Y}$) and no mutational correlation.

##### Set 2: Selection, genetic drift, and unequal mutation

These simulations are identical to Set 1 except that we varied $V_{M,Y}$ over $1{\times}10^{-4}$, $5{\times}10^{-4}$, 0.002, 0.01 while keeping $V_{M,X} = 0.001$. These simulations test whether unequal mutational variance in constituent traits could shape their covariance.

##### Set 3: Selection, genetic drift, and mutational correlation

These simulations are identical to Set 1 except that we varied $\rho_{M,XY}$ over $-0.9, -0.45, 0.45, 0.9$. These simulations test whether mutational covariance in constituent traits could shape their covariance.

##### Set 4: Selection, genetic drift, and physical limits

These simulations are conceptually similar to Set 1 except that we applied physical limits on the fraction of epidermal area allocated to stomata ($f_S$) and therefore values had to be on the measurement scale. When $\beta = 0.5$, $z_0 = \log(4.8 \times 10^{-4}$ mol m$^{-2}$ s$^{-1})$; when $\beta = 1.0$, $z_0 = -\log(10^6)$ to convert stomatal density from mm$^{-2}$ to $\mu$m$^{-2}$.

We also used realistic values for the long-run average. When $\beta = 0.5$, $\theta = \log(1.32$ mol m$^{-2}$ s$^{-1})$, the average $\log(g_{\max})$ among species in our data; when $\beta = 1.0$, $\theta = \log(0.044)$, the average $\log(f_S)$ among species in our data. To be consistent with other simulations, we set macroevolutionary parameters so that the stationary distribution of $z$ had variance ($\sigma^2/(2\alpha) = 0.5$), but changed $\sigma = \sqrt{0.001}$ and $\alpha = 5{\times}10^{-6}$ to keep the ratio between variance components ($\sigma^2$, $V_E$, $V_M$, and $V_S$) consistent across simulation sets.

We set $V_E = 0.005$ to be 1% of the stationary variance in $z$ so that intraspecific variation was small relative to interspecific variance, as we expect in phylogenetically broad comparative studies. As in Set 1, $1000V_M = V_E$ and $V_S = 10V_E$. Mutational variance in constituent traits were equal and we assumed no mutational covariance.

Whereas in previous simulations there was no limit on trait values, we applied a physical limit such that $f_S$ could not exceed 0.337, the maximum value in our data set. Higher values such as 0.4 or 0.5 did not qualitatively alter results (results not shown). To impose physical limitations, we treated the maximum $f_S$ as a reflecting boundary for any environmental or mutational deviations that caused $f_S > f_{S,\max}$. In the $x, y$ plane, the physical limit is described by a line $y = f_{S,\max} - x$. Environmental or mutational deviations can be described as a vector in the $x, y$ plane with a corresponding direction and magnitude. If the trait vector hits the physical limit, it reflects off following the law of reflection (incident angle = reflected angle) and the total magnitude is preserved to determine the final phenotype after environmental or mutational effects. While reflecting boundaries alter the trait covariance when species are near it, they are mathematically preferable to imposing limits by altering the magnitude of mutational changes near the physical limit as this would cause an alternative equilibrium state where all species eventually get "stuck" near the physical limit, which is unrealistic.

Table S4: Parameter values for simulations. Each parameter has a mathematical symbol used in the text, the $R$ character string used in the source code, a brief description, and the value(s) used for each simulation set. The text explains how we chose appropriate parameter values.

| Symbol | $R$ character | Description | Set 1 | Set 2 | Set 3 | Set 4 |
|---|---|---|---|---|---|---|
| **Trait parameters:** | | | | | | |
| $\beta$ | `beta` | scaling coefficient of composite trait equation | 0.5, 1 | 0.5, 1 | 0.5, 1 | 0.5, 1 |
| $z_0$ | `z_0` | intercept of composite trait equation | 0 | 0 | 0 | $\log(4.8 \times 10^{-4}),\ -\log(10^6)$ |
| **Macroevolutionary adaptive landscape parameters:** | | | | | | |
| $\alpha$ | `alpha` | return rate toward $\theta$ | 0.001 | 0.001 | 0.001 | $5{\times}10^{-6}$ |
| $\sigma$ | `sigma` | rate of random movement in $z_{\text{opt}}$ | $\sqrt{0.001}$ | $\sqrt{0.001}$ | $\sqrt{0.001}$ | $\sqrt{5 \times 10^{-6}}$ |
| $\theta$ | `theta` | long-run average $z_{\text{opt}}$ | 0 | 0 | 0 | $\log(1.32),\ \log(0.044)$ |
| **Microevolutionary quantitative genetic parameters:** | | | | | | |
| $N$ | `N` | population size | 1024 | 1024 | 1024 | 1024 |
| $\rho_{M,XY}$ | `r_XY` | mutational correlation between constituent traits | 0 | 0 | -0.9, -0.45, 0.45, 0.9 | 0 |
| $V_E$ | `V_e` | environmental variance in constituent traits | 1 | 1 | 1 | 0.005 |
| $V_{M,X}$ | `V_mX` | mutational variance in constituent trait $X$ | 0.001 | 0.001 | 0.001 | $5{\times}10^{-6}$ |
| $V_{M,Y}$ | `V_mY` | mutational variance in constituent trait $Y$ | 0.001 | $1{\times}10^{-4}$, $5{\times}10^{-4}$, 0.002, 0.01 | 0.001 | $5{\times}10^{-6}$ |
| $V_S$ | `V_s` | strength of stabilizing selection on $z$ | 10 | 10 | 10 | 0.05 |
| $W_{\max}$ | `W_max` | Maximum fitness when $z = z_{\text{opt}}$ | 1 | 1 | 1 | 1 |
| **Physical limits:** | | | | | | |
| $f_{S,\max}$ | `max_fs` | Maximum possible $f_S$ | $\infty$ | $\infty$ | $\infty$ | 0.337 |

### Simulation results

In all simulations sets, we estimated $\hat{\beta}$ and 99% confidence intervals using standard linear regression methods with the `lm()` function in R version 4.0.5. If our original proposition is correct, then we expect scaling estimates ($\hat{\beta}$) to recover simulated values. If primary constraint occurs on $g_{\text{max}}$ (stomatal-area increase hypothesis) then we will estimate $\hat{\beta} = 0.5$; when primary constraint occurs on $f_S$ (stomatal-area minimization hypothesis) then we will estimate $\hat{\beta} = 1.0$.

In all simulation sets, we observed that the linear regression slope, multiplied by -1, is an unbiased or nearly unbiased estimator of $\beta$. In Set 1, which included genetic drift, assumed equal mutational variance in constituent traits and no mutational correlation, we found that simulated and estimated values of $\beta$ matched (Figure S2). Similarly, in Set 2, we found that even 10-fold differences in the mutational variance of constituent traits had no effect (Figure S3). In Set 3, our estimates slightly underestimated $\beta$ when the the mutational correlation between constituent traits ($\rho_{M,XY} = 0.9$) is very strongly orthogonal to the negative among-species covariance (Figure S4). Even in this situation, the bias is small compared to the difference in predicted values between competing hypotheses. Similarly, in Set 4, physical limits on the maximum stomatal cover led to slight underestimates, but the bias is small compared to difference in predicted values (Figure S5). The confidence intervals are wider because imposing a maximum stomatal cover reduced the variance in constituent traits, while variance in the composite trait remained the same at stationarity. This means that interspecific variation is spread further around the main axis of variation rather than along the main axis of variation, leading to greater uncertainty in the parameter estimate.

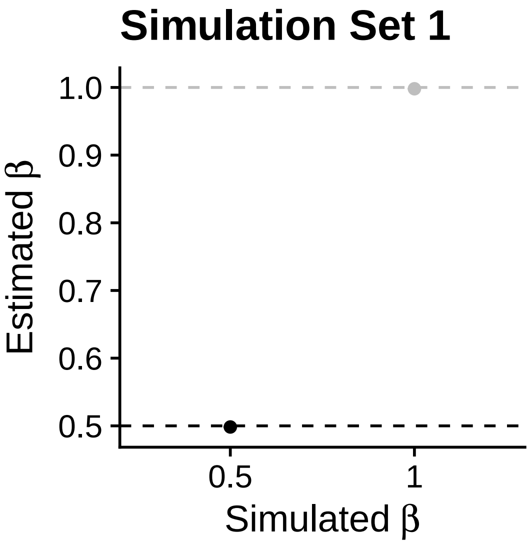


Figure S2: Forward-time, macroevolutionary quantitative genetic simulations demonstrate that the linear regression slope accurately estimates the simulated scaling values ("Simulated $\beta$") across species. When $\beta$ is simulated to be 0.5 (stomatal-area increase hypothesis) we expect the estimate $\hat{\beta} = 0.5$ (black dashed line and point); when $\beta$ is simulated to be 1.0 (stomatal-area minimization hypothesis) we expect the estimate $\hat{\beta} = 1.0$ (grey dashed line and point). The 99% confidence intervals are too narrow to see because they do not extend beyond the size of the point estimate. See Table S4 for information on simulation parameters.

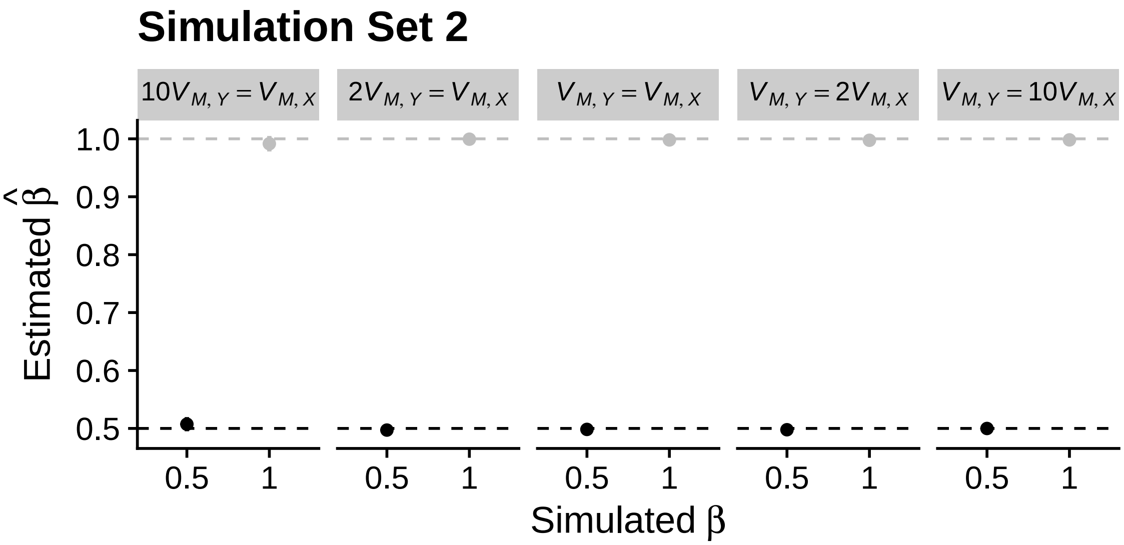


Figure S3: Forward-time, macroevolutionary quantitative genetic simulations demonstrate that the linear regression slope accurately estimates the simulated scaling values ("Simulated $\beta$") across species. When $\beta$ is simulated to be 0.5 (stomatal-area increase hypothesis) we expect the estimate $\hat{\beta} = 0.5$ (black dashed lines and points); when $\beta$ is simulated to be 1.0 (stomatal-area minimization hypothesis) we expect the estimate $\hat{\beta} = 1.0$ (grey dashed lines and points). The panels, from left to right, display the results for simulations where the mutational variance in $Y$ was 10× lower than that $X$ to where it was 10× higher than that $X$. The 99% confidence intervals are too narrow to see because they do not extend beyond the size of the point estimate. See Table S4 for information on simulation parameters.

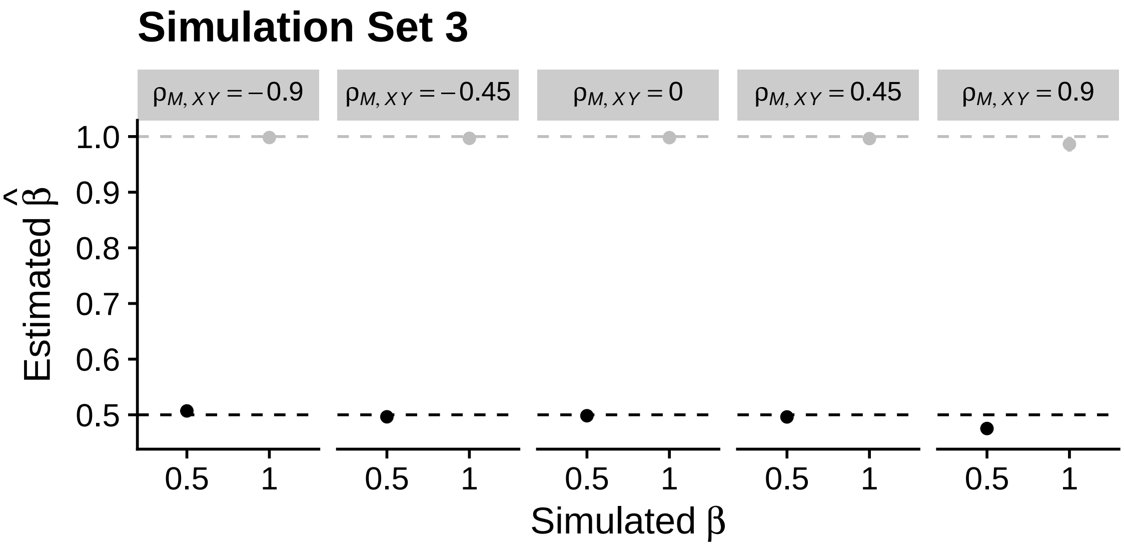


Figure S4: Forward-time, macroevolutionary quantitative genetic simulations demonstrate that the linear regression slope accurately estimates the simulated scaling values ("Simulated $\beta$") across species. When $\beta$ is simulated to be 0.5 (stomatal-area increase hypothesis) we expect the estimate $\hat{\beta} = 0.5$ (black dashed lines and points); when $\beta$ is simulated to be 1.0 (stomatal-area minimization hypothesis) we expect the estimate $\hat{\beta} = 1.0$ (grey dashed lines and points). The panels, from left to right, display the results for simulations where the mutational correlation between $X$ and $Y$ increases from $\rho_{M,XY} = -0.9$ to $\rho_{M,XY} = 0.9$. The 99% confidence intervals are too narrow to see because they do not extend beyond the size of the point estimate. See Table S4 for information on simulation parameters.

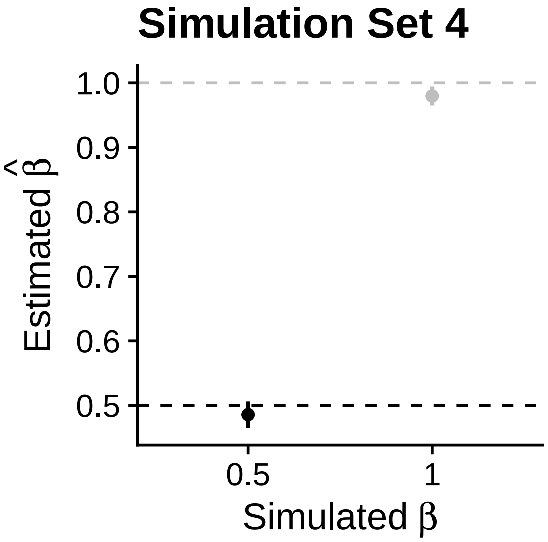


Figure S5: Forward-time, macroevolutionary quantitative genetic simulations demonstrate that the linear regression slope accurately estimates the simulated scaling values ("Simulated $\beta$") across species. When $\beta$ is simulated to be 0.5 (stomatal-area increase hypothesis) we expect the estimate $\hat{\beta} = 0.5$ (black dashed line and point); when $\beta$ is simulated to be 1.0 (stomatal-area minimization hypothesis) we expect the estimate $\hat{\beta} = 1.0$ (grey dashed line and point). The error bars indicate the 99% confidence intervals. These simulations are similar to Simulation Set 1 except that stomatal cover cannot exceed a maximum value. See Table S4 for information on simulation parameters.
