## Supplemental Material 3 for "Scaling between stomatal size and density in forest plants" (from another version of this preprint: version 1, 2021-04-26)

### Supplementary Information: Additional R packages

| package | version | reference |
|---|---|---|
| adaptMCMC | 1.3 | Scheidegger & and [1] |
| ape | 5.4-1 | Paradis & Schliep [2] |
| arrayhelpers | 1.1-0 | Beleites [3] |
| assertthat | 0.2.1 | Wickham [4] |
| backports | 1.2.1 | Lang & R Core Team [5] |
| broom | 0.7.3 | Robinson *et al.* [6] |
| callr | 3.5.1 | Csárdi & Chang [7] |
| cellranger | 1.1.0 | Bryan [8] |
| checkmate | 2.0.0 | Lang [9] |
| cli | 2.2.0 | Csárdi [10] |
| coda | 0.19-4 | Plummer *et al.* [11] |
| codetools | 0.2-18 | Tierney [12] |
| colorspace | 2.0-0 | Zeileis *et al.* [13] |
| | | Zeileis *et al.* [14] |
| | | Stauffer *et al.* [15] |
| cowplot | 1.1.0 | Wilke [16] |
| crayon | 1.3.4 | Csárdi [17] |
| data.table | 1.13.4 | Dowle & Srinivasan [18] |
| DBI | 1.1.0 | R Special Interest Group on Databases (R-SIG-DB) *et al.* [19] |
| dbplyr | 2.0.0 | Wickham & Ruiz [20] |
| desc | 1.2.0 | Csárdi *et al.* [21] |
| devtools | 2.3.2 | Wickham *et al.* [22] |
| digest | 0.6.27 | **lucas2020digest** |
| distributional | 0.2.1 | O'Hara-Wild & Hayes [23] |
| dplyr | 1.0.2 | Wickham *et al.* [24] |

| package | version | reference |
|---|---|---|
| ellipse | 0.4.2 | Murdoch & Chow [25] |
| ellipsis | 0.3.1 | Wickham [26] |
| evaluate | 0.14 | Wickham & Xie [27] |
| fansi | 0.4.1 | Gaslam [28] |
| farver | 2.0.3 | Pedersen *et al.* [29] |
| filehash | 2.4-2 | Peng [30] |
| forcats | 0.5.0 | Wickham [31] |
| foreach | 1.5.1 | Microsoft & Weston [32] |
| fs | 1.5.0 | Hester & Wickham [33] |
| furrr | 0.2.1 | Vaughan & Dancho [34] |
| future | 1.21.0 | Bengtsson [35] |
| future.apply | 1.6.0 | Bengtsson [36] |
| generics | 0.1.0 | Wickham *et al.* [37] |
| ggdist | 2.3.0 | Kay [38] |
| ggplot2 | 3.3.2 | Wickham [39] |
| globals | 0.14.0 | Bengtsson [40] |
| glue | 1.4.2 | Hester [41] |
| gtable | 0.3.0 | Wickham & Pedersen [42] |
| haven | 2.3.1 | Wickham & Miller [43] |
| hms | 0.5.3 | Müller [44] |
| htmltools | 0.5.0 | Cheng *et al.* [45] |
| httr | 1.4.2 | Wickham [46] |
| iterators | 1.0.13 | Analytics & Weston [47] |
| jsonlite | 1.7.2 | Ooms [48] |
| knitr | 1.30 | Xie [49] |
| | | Xie [50] |
| | | Xie [51] |
| lamW | 1.3.3 | Adler [52] |
| lattice | 0.20-41 | Sarkar [53] |
| lifecycle | 0.2.0 | Henry [54] |
| listenv | 0.8.0 | Bengtsson [55] |

| package | version | reference |
|---|---|---|
| lubridate | 1.7.9.2 | Grolemund & Wickham [56] |
| magrittr | 2.0.1 | Bache & Wickham [57] |
| Matrix | 1.3-2 | Bates & Maechler [58] |
| memoise | 1.1.0 | Wickham *et al.* [59] |
| modelr | 0.1.8 | Wickham [60] |
| munsell | 0.5.0 | Wickham [61] |
| mvnfast | 0.2.5.1 | Fasiolo [62] |
| nlme | 3.1-152 | Pinheiro *et al.* [63] |
| parallelly | 1.22.0 | Bengtsson [64] |
| phylolm | 2.6.2 | Ho & Ane [65] |
| pillar | 1.4.7 | Müller & Wickham [66] |
| pkgbuild | 1.2.0 | Wickham & Hester [67] |
| pkgconfig | 2.0.3 | Csárdi [68] |
| pkgload | 1.1.0 | Wickham *et al.* [69] |
| plyr | 1.8.6 | Wickham [70] |
| POUMM | 2.1.7 | Mitov & Stadler [71] |
| | | Mitov & Stadler [72] |
| prettyunits | 1.1.1 | Csardi [73] |
| processx | 3.4.5 | Csárdi & Chang [74] |
| progress | 1.2.2 | Csárdi & FitzJohn [75] |
| ps | 1.5.0 | Loden *et al.* [76] |
| purrr | 0.3.4 | Henry & Wickham [77] |
| R6 | 2.5.0 | Chang [78] |
| Rcpp | 1.0.5 | Eddelbuettel & François [79] |
| | | Eddelbuettel [80] |
| | | Eddelbuettel & Balamuta [81] |
| RcppParallel | 5.0.2 | Allaire *et al.* [82] |
| readr | 1.4.0 | Wickham & Hester [83] |
| readxl | 1.3.1 | Wickham & Bryan [84] |
| RefManageR | 1.3.0 | McLean [85] |
| | | McLean [86] |

| package | version | reference |
|---|---|---|
| remotes | 2.2.0 | Hester *et al.* [87] |
| reprex | 0.3.0 | Bryan *et al.* [88] |
| rlang | 0.4.9 | Henry & Wickham [89] |
| rmarkdown | 2.7 | Allaire *et al.* [90] |
| | | Xie *et al.* [91] |
| | | Xie *et al.* [92] |
| rprojroot | 2.0.2 | Müller [93] |
| rstudioapi | 0.13 | Ushey *et al.* [94] |
| rvest | 0.3.6 | Wickham [95] |
| scales | 1.1.1 | Wickham & Seidel [96] |
| sessioninfo | 1.1.1 | Csárdi *et al.* [97] |
| stringi | 1.5.3 | Gagolewski [98] |
| stringr | 1.4.0 | Wickham [99] |
| svUnit | 1.0.3 | Grosjean [100] |
| taxonlookup | 1.1.5 | Cornwell *et al.* [101] |
| testthat | 3.0.1 | Wickham [102] |
| tibble | 3.0.4 | Müller & Wickham [103] |
| tidybayes | 2.3.1 | Kay [104] |
| tidyr | 1.1.2 | Wickham [105] |
| tidyselect | 1.1.0 | Henry & Wickham [106] |
| tidyverse | 1.3.0 | Wickham *et al.* [107] |
| tikzDevice | 0.12.3.1 | Sharpsteen & Bracken [108] |
| units | 0.6-7 | Pebesma *et al.* [109] |
| usethis | 2.0.0 | Wickham & Bryan [110] |
| vctrs | 0.3.6 | Wickham *et al.* [111] |
| withr | 2.3.0 | Hester *et al.* [112] |
| xfun | 0.19 | Xie [113] |
| xml2 | 1.3.2 | Wickham *et al.* [114] |
| yaml | 2.2.1 | Stephens *et al.* [115] |
